# Palmitate induces apoptotic cell death and inflammasome activation in human placental macrophages

**DOI:** 10.1101/799718

**Authors:** Lisa M. Rogers, Carlos H. Serezani, Alison J. Eastman, Alyssa H. Hasty, Linda Englund-Ögge, Bo Jacobsson, Kasey C. Vickers, David M. Aronoff

## Abstract

**Introduction:** There is an increasing prevalence of non-communicable diseases worldwide. Metabolic diseases such as obesity and gestational diabetes mellitus (GDM) increasingly affect women during pregnancy, which can harm pregnancy outcomes and the long-term health and wellbeing of exposed offspring. Both obesity and GDM have been associated with proinflammatory effects within the placenta, the critical organ governing fetal development.

**Methods:** The purpose of these studies was to model, *in vitro*, the effects of metabolic stress (high levels of glucose, insulin and saturated lipids) on placental macrophage biology, since these cells are the primary innate immune phagocyte within the placenta with roles in governing maternofetal immune tolerance and antimicrobial host defense. Macrophages were isolated from the villous core of term, human placentae delivered through nonlaboring, elective Cesarean sections and exposed to combinations of elevated glucose (30 mM), insulin (10 nM) and the saturated lipid palmitic acid (palmitate, 0.4 mM).

**Results:** We found that palmitate alone induced the activation of the nucleotide-binding oligomerization domain-like receptor (NLR) Family Pyrin Domain Containing 3 (NLRP3) inflammasome in placental macrophages, which was associated with increased interleukin 1 beta release and an increase in apoptotic cell death. Glucose and insulin neither provoked these effects nor augmented the impact of palmitate itself.

**Discussion:** Our findings confirm an impact of saturated fat on placental macrophage immune activation and could be relevant to the impact of metabolic stress *in vivo*.

## Introduction

Modern civilization is experiencing a shift towards non-communicable diseases and away from infectious disease [1]. Metabolic diseases such as obesity and type 2 diabetes mellitus are non-communicable diseases that increasingly affect women during pregnancy, which can have detrimental effects on pregnancy outcomes, long-term health of the woman and also negative long-term health effects on exposed offspring. Worldwide, more than 20 million births each year are impacted by some form of hyperglycemia in pregnancy, including gestational diabetes mellitus (GDM) or preexisting diabetes [1]. In addition, maternal prepregnancy obesity is a risk factor for early life overweight and obesity in offspring [2]. Prepregnancy obesity greatly increases the risk for GDM, closely linking these two disorders [3]. The combination of maternal obesity and diabetes, so called diabesity syndrome [4, 5], is particularly important in conferring risk for metabolic diseases to the next generation [3, 6, 7].

How obesity and diabetes impact fetal development and the risk for long term health outcomes for offspring remains incompletely addressed, though a combination of cellular/tissue damage and/or epigenetic modification appear to be involved [8]. The placenta is a critical organ orchestrating nutrient transport to the fetus, hormone production, maternofetal gas exchange, removal of waste, maintenance of maternofetal immune tolerance, and host defense. It is thus an important biological conduit to mediate the non-genomic transmission of risk for non-communicable diseases [9, 10]. Both obesity and diabetes are associated with low-grade increases in inflammation throughout the body [3, 11, 12]. Placental inflammation has been observed in pregnancies complicated by obesity [13] and GDM [14], and has been suggested to play a central role in determining the fetal environment in such pregnancies [3]. Recent research in a mouse model of pregnancy-associated diabesity revealed alterations in the expression of genes governing immunity and inflammation within the placenta [15]. How metabolic stressors such as obesity and/or diabetes (or both) modulate gene expression in the placenta remains to be defined. It has been suggested that circulating, saturated free fatty acids (FFAs), such as the ω-6 FFA palmitate, can provoke inflammatory responses in placental cells such as trophoblasts by activating the NLRP3 inflammasome [16, 17]. This might be relevant because palmitate levels are elevated in the systemic circulation of pregnant women with prepregnancy obesity [18] and GDM [19]. In addition, lipid levels are higher in placental tissues from obese compared to lean women and trophoblasts obtained from obese women more rapidly esterify palmitate compared to cells obtained from lean women [20]. The extent to which either high glucose or insulin also contribute to placental inflammation in the setting of diabesity (or GDM in the absence of obesity) is unclear.

Macrophages within the placenta are important innate immune sentinels capable of sensing the environment and producing proinflammatory mediators in response to stress. Tissue-based studies suggest that metabolic stress alters macrophage abundance in the placenta [21, 22]. However, the exact contribution of macrophages to placental inflammatory tone in the face of obesity or diabetes has not been clearly defined. Thus, in the present set of studies, we addressed the hypothesis that modeling diabesity *in vitro* with isolated human placental macrophages would provoke a proinflammatory response that reflects combined effects of exposure to excessive glucose, insulin and saturated FFA (palmitate) in the *in vitro*.

## Methods

### Reagents

RPMI 1640 culture medium (glucose-containing & glucose-free), antibiotic solution (penicillin and streptomycin) including an antimycotic (amphotericin), Alexa Fluor 488 annexin V/dead cell apoptosis kit, cell extraction buffer, Trizol Reagent, human active caspase-3 ELISA kit, TaqMan Gene Expression Assay Primers (FAM; IL-1β & GAPDH), and phosphate buffered saline (PBS) were all purchased from Invitrogen (Carlsbad, CA). Paraformaldehyde was purchased from Alfa Aesar (Ward Hill, MA). Charcoal-stripped and dextran-treated fetal bovine serum (FBS) was purchased from HyClone Laboratories (Waltham, MA). Protease inhibitor cocktail, deoxyribonuclease from bovine pancreas (type IV), hyaluronidase from bovine testes (type I-S), collagenase from C. histolyticum (type I-A), percoll, glucose, Duolink In Situ PLA Kit, Phenylmethylsulfonyl fluoride (PMSF), and sodium bicarbonate were from Sigma-Aldrich (St. Louis, MO). Lysis buffer for erythrocytes and human CD68-PE (clone Y1/82A) were from eBiosciences (San Diego, CA). Permeabilization buffers for flow cytometry were purchased from BD Biosciences (San Diego, CA). Human CD14 microbeads and LS columns were purchased from Miltenyi Biotec (San Diego, CA). Optiblot Bradford Reagent was from Abcam (Cambridge, MA). Caspase-1 (p20 subunit), IL-1β, TNF-α, IL-6, and CCL2 ELISA kits and all supplementary reagents were from R&D Systems (Minneapolis, MN). Novolin human insulin was purchased from Novo Nordisk (Plainsboro, NJ). Palmitic acid (palmitate, C16:O) was obtained from Nu-Chek Prep (Elysian, MN). Anti-ASC pAB (AL177) and anti-NLRP3 mAB (Cryo-2) were purchased from AdipoGen (San Diego, CA). ProLong Gold Antifade with DAPI was from Cell Signaling Technologies (Danvers, MA). The irreversible caspase-1 inhibitor VI (Z-YVAD-FMK) was from Millipore (obtained through Thermo-Fisher). iQ Supermix, iScript cDNA synthesis kit, and iScript Reverse Transcription Supermix for RT-qPCR were from Bio-Rad Laboratories (Hercules, California).

### Human Subjects

Placental tissue samples were provided by the Cooperative Human Tissue Network which is funded by the National Cancer Institute. All tissues were collected in accordance with Vanderbilt University Institutional Review Board (approval #131607 & 181998) and Declaration of Helsinki. Recruitment was limited to term, non-laboring women where the placentae were obtained at the time of Cesarean section. Samples were only obtained from healthy donors with no significant medical condition, aged 18-40 years. Because samples were deidentified prior to research use, data were not available regarding maternal or neonatal covariates.

### Placental Macrophage Isolation

Human placental macrophages from the villous core were isolated as previously described [23]. Briefly, placental cotyledon tissue, separated from the maternal decidua basalis, was vigorously washed with PBS to remove circulating blood. The tissue was minced into small pieces and weighed to determine final grams collected. Tissue fragments were placed into 250 mL sterile bottles with sterile digestion solution containing 150 μg/mL deoxyribonuclease, 1 mg/mL collagenase, and 1 mg/mL hyaluronidase at 10 mL per gram of tissue. Placenta was digested for 1 h at 37°C, shaking at 180 RPM. Digested tissue was filtered through a 280 μm metal sieve, followed by 180 and 80 μm nylon screens. Cells were centrifuged at 1500 RPM and resuspended in 25% Percoll diluted in cold RPMI media containing 10% FBS and 1% antibiotic-antimycotic (referred hereafter as RPMI^+/+^) and overlaid onto 50% Percoll, plus 2 mL of PBS on top of the density gradient. CD14^+^ macrophages were isolated by positive selection using the magnetic MACS^®^ large cell separation column system according to the manufacturer’s instructions. Isolated CD14^+^ placental macrophages were plated and rested overnight in RPMI^+/+^ at 37°C with 5% CO2 before experimentation. Purity of PM samples was determined to be 98.04 ± 0.58 % (mean ± SEM) based on CD68 positivity (data not shown).

### Metabolic Stress Treatment; Cytokine and Lysate Analysis

Placental macrophages were cultured at a concentration of 3 × 10^6^ cells/mL in RPMI^+/+^ and rested overnight after isolation. Macrophages were then acclimated to glucose-free RPMI^+/+^ for 60 min (in the presence or absence of the irreversible caspase-1 inhibitor VI (Z-YVAD-FMK; 10 μM)) before any glucose treatment was performed. All glucose treatments were performed in RPMI^+/+^. Placental macrophages were exposed to either 5 mM glucose (euglycemia; control media), 30 mM glucose (a model of hyperglycemia), 30 mM glucose + 0.4 mM palmitate + 10 nM human insulin (together known as a metabolic cocktail, MetaC, a model of diabesity [24]), 5 mM glucose + 0.4 mM palmitate (Palm), or 5 mM glucose + 10 nM human insulin for time points between 1 and 24 h. For a dose curve at 24 h, we assessed palmitate doses between 0 and 0.8 mM in the presence of 5 mM glucose. Supernatants were harvested for cytokine analysis by ELISA, following the manufacturers instruction for each analyte (IL-1β, Caspase-1 p20). Cellular lysates were harvested for active caspase-3 quantification.

### Inflammasome proximity ligation assay (PLA)

Placental macrophages were assessed for inflammasome complex interactions using the Duolink PLA immunofluorescence assay format. Briefly, 0.3 × 10^6^ placental macrophages were cultured in chamber slides and treated for 4 h with glucose or the metabolic stress components as they were for cytokine analysis above. Cells were then washed with PBS, fixed and permeabilized, blocked with kit-supplied blocking buffer, and stained with primary antibodies anti-NLRP3 & anti-ASC (1:200) in kit-included antibody diluent. Placental macrophages were then washed and incubated with the necessary probes before ligation and amplification took place. Final washes were completed, and chamber slides were cover-slipped with Antifade Prolong Gold with DAPI and let dry overnight in the dark before imaging (Nikon Eclipse fluorescent microscope) and quantification with Image-J software.

### Flow Cytometry Cellular Death Analysis

Placental macrophages were plated, rested, and exposed to glucose or the metabolic stress components as they were for cytokine analysis above. Treatments were performed for 4 or 24 h before cells were harvested for flow cytometric analysis. Cellular death was assayed by dual-staining with annexin V-AF 488 and propidium iodide (PI) following manufacturer’s instructions. Briefly, some PMs were stained with anti-human CD68 to assess purity before being stained with the annexin/PI staining kit. Paraformaldehyde-fixed placental macrophages were acquired immediately after staining on an LSRII 5-laser flow cytometer (BD Biosciences). Living cells were defined as PI-/annexin-; necrotic cells were defined as PI+/annexin-; early apoptotic cells were defined as PI-/annexin+; and late apoptotic or pyroptotic cells were defined as PI+/annexin+. Data files were analyzed using BD Diva (BD Biosciences).

### Quantitative Real Time (RT)-PCR

Placental macrophages were plated, rested, and exposed to glucose or the metabolic stress components for 4 h as they were for cytokine analysis above. Total RNA was isolated using TRIzol reagent according to the manufacturer’s instructions. Template RNA (325 ng) was reverse transcribed using the iScript cDNA Synthesis Kit and cDNA was amplified for IL-1β and GAPDH under the following thermal cycling conditions: 3 m at 95°C, 12 s at 95°C, and 45 s at 60°C for 40 cycles. Data are quantified and presented using the 2^−Δct^ method.

### Statistical Analysis

All experiments were performed in multiple, independent biological replicates as noted in each figure. Data points from independent experiments are presented along with means and analyzed with GraphPad Prism 6.0 software (GraphPad Software, San Diego, CA). Comparisons between two experimental groups were performed with a paired Student *t* test, while comparisons between three or more groups were analyzed with a repeated measures 1-way ANOVA with a Dunnett or Tukey multiple comparisons post-test where appropriate. Differences were considered statistically significant for *P <0.05*.

## Results

### Palmitate increases the release of proinflammatory cytokine IL-1β and cleaved caspase-1 in placental macrophages

Human placental macrophages were exposed *in vitro* to models of hyperglycemia, hyperinsulinemia and saturated fat hyperlipidemia, or to a combination of these factors (MetaC; a model of diabesity [24]) as detailed above, for durations ranging from 1 to 24 hours. These experiments revealed a time-dependent increase in both secreted IL-1β and cleaved caspase-1 (p20 subunit) in response to MetaC or palmitic acid (**FIG 1**). By 1 hour, there were no changes in IL-1β among any groups (**Fig. 1A-B**), while caspase-1 was slightly but significantly increased in the MetaC condition relative to control; palmitate treatment did not alter caspase-1 levels by 1 hour (**Fig. 1G-H**). By 4 hours, a significant increase in MetaC-induced IL-1β and cleaved caspase-1 was observed relative to the euglycemic control (**FIG 1C-D, I-J**), differences that were further increased at 24 hours (**FIG 1E-F, K-L**). Exposure of placental macrophages to hyperinsulinemia alone or hyperglycemia alone for 24 hours did not result in increased secretion of IL-1β or cleaved caspase-1 (**SUPPL FIG 1**), suggesting palmitate alone is responsible for the secreted IL-1β and cleaved caspase-1 we are detecting. Exposing placental macrophages to a range of concentrations of palmitate (range of 0 - 0.8 mM) for 24 hours yielded a dose-dependent increase in both secreted IL-1β and cleaved caspase-1 (p20 subunit) (**SUPPL FIG 2**). The irreversible caspase-1 inhibitor VI (Z-YVAD-FMK) applied for 1 hour before 5 mM glucose (euglycemia) or MetaC treatment reduced secreted IL-1β at both 4 hours and 24 hours (**FIG 2A-B**). We did not observe increases in other proinflammatory cytokines such as TNFα, CCL2, or IL-6 in response to MetaC (**SUPPL FIG 5B-D**).

**Figure 1:**
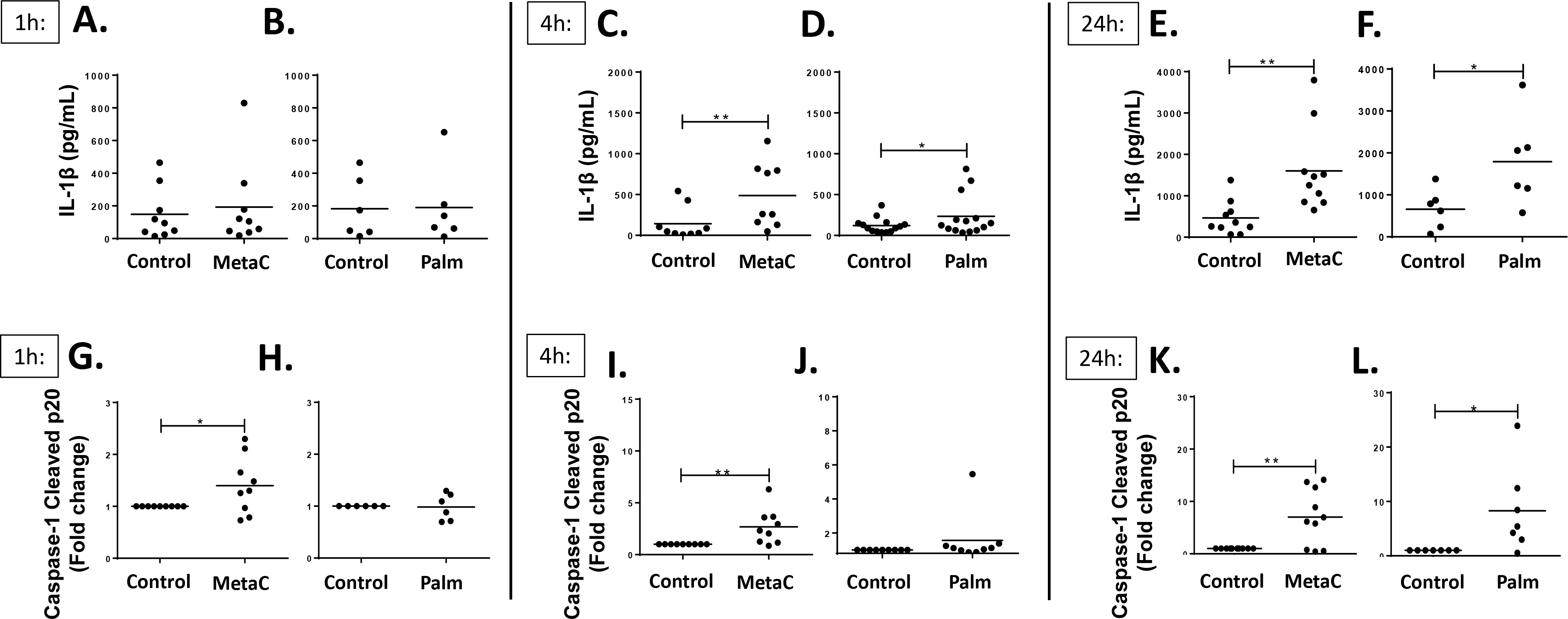
Metabolic stress increases the release of proinflammatory cytokine IL-1β via caspase-1 in placental macrophages in a time-dependent manner. Placental macrophages were treated for 1 - 24 h with 5 mM glucose (control), MetaC, or palmitate (0.4 mM) before supernatant was harvested for IL-1β **(A-F)** (n=3-9) and cleaved caspase-1 **(G-L)** (n=3-9) ELISA analysis. Paired Student *t*-test; mean ± SEM for all graphs; *\*p<0.05* and *\*\*p<0.01*.

**Figure 2:**
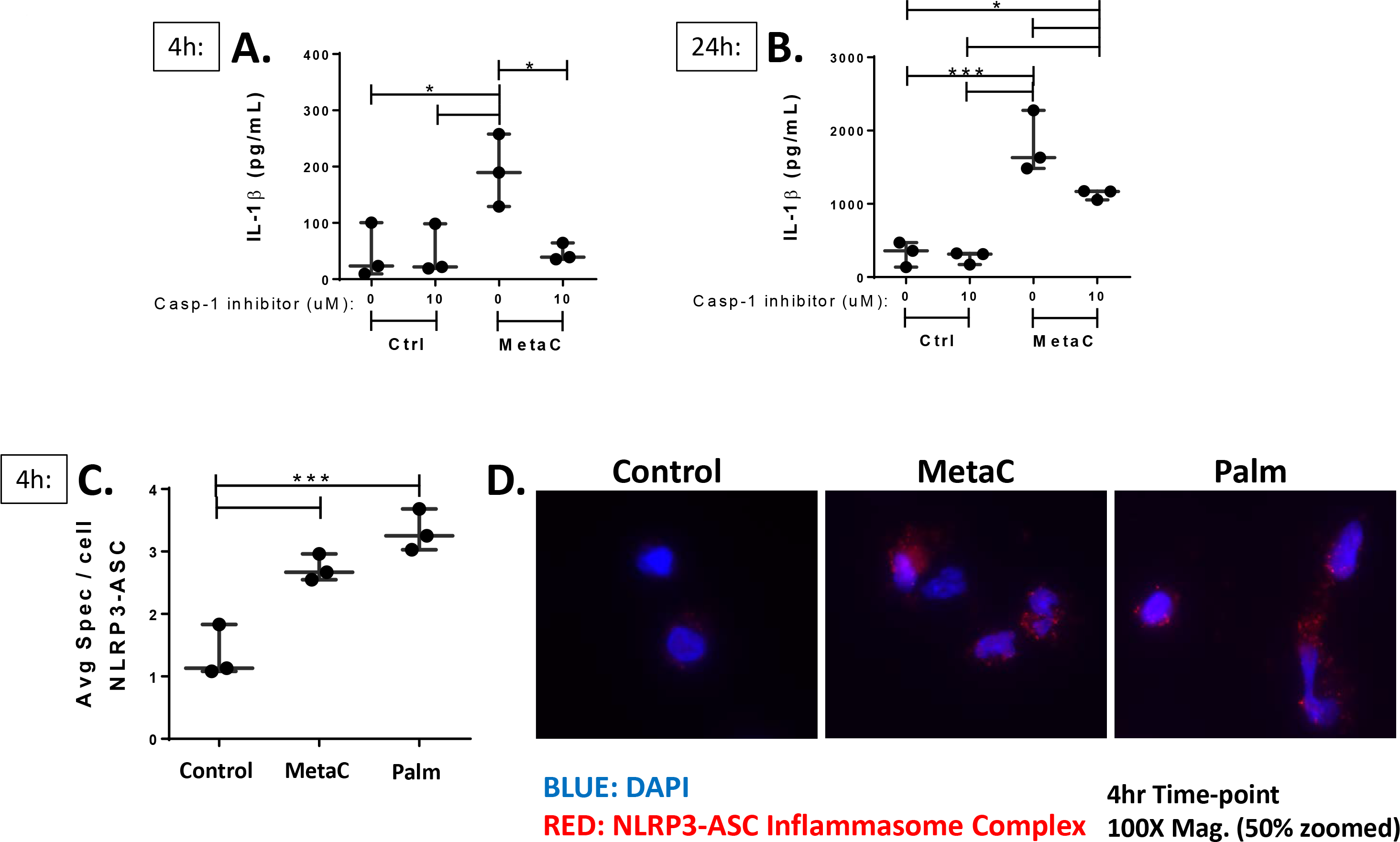
Metabolic stress-induced secretion of IL-1β by placental macrophages is blocked by caspase-1 inhibition; metabolic stress activates assembly of the NLRP3-ASC inflammasome complex. Placental macrophages (PM) were pretreated for 1 h with an irreversible caspase-1 inhibitor before being exposed to 5 mM glucose or MetaC. Supernatants were harvested for IL-1β ELISA analysis at 4 h **(A)** and 24 h **(B)** (n=3 each). Paired Student *t*-test; mean ± SEM, *p*<0.05* and \*\*\**p<0.001*. PM cells were treated for 4 hours with 5 mM glucose, MetaC, or palmitate. Cells were stained using the proximal ligation assay for assembly of the NLRP3 and ASC inflammasome complex. **(C)** Average spec/cell quantification by automatic particle counting using Image-J software (n=3). 1-way repeated measures ANOVA with Tukey post-test; mean ± SEM; \*\*\**p<0.001*. **(D)** Representative fluorescent images of treated macrophages (100X; blue represents NucBlue Live (Hoechst 33342) and red represents NLRP3-ASC inflammasome complex).

### Metabolic stress activates assembly of the NLRP3-ASC inflammasome complex

The inflammasome promotes the maturation and secretion of proinflammatory IL-1β [25]. Human placental macrophages were treated with 5 mM glucose (euglycemia), MetaC, or palmitate for 4 hours and assembly of the NLRP3-ASC inflammasome complex was tracked by PLA as described in the *Methods* section. MetaC and palmitate resulted in significantly more assembled NLRP3-ASC complexes per cell compared to euglycemic placental macrophages (**FIG 2C-D**).

### Metabolic stress induces multiple types of cellular death

Human placental macrophages were treated with 5 mM glucose (euglycemia), MetaC, palmitate, 30 mM glucose (hyperglycemia) or insulin alone (hyperinsulinemia) for 4 or 24 hours and assayed for cell death by flow cytometry using annexin V staining, which detects apoptotic changes, and PI staining for necrosis [26]. An example of gating methods are shown (**FIG 3A**). MetaC and palmitate treatment for 4 h induced significant increases in multiple types of cellular death: necrosis (PI+/annexin−), early apoptosis (PI−/annexin+), and late apoptosis and/or pyroptosis (PI+/annexin+) (**FIG 3B**, **SUPPL FIG 3**) compared to euglycemia. Early apoptotic cell death was exacerbated by 24 h of incubation with palmitate, resulting in 43.14 ± 9.13 % (mean ± SEM) of cells undergoing such apoptosis (**FIG 3C**, **SUPPL FIG 4**). Hyperglycemia or insulin alone did not differ from euglycemia. The majority of MetaC-induced and palmitate-induced cellular death detected was early apoptosis. We also investigated the presence of cleaved caspase-3, an important marker of the cell’s entry point into the apoptotic signaling pathway [27] and found a significant increase in cleaved caspase-3 in the cellular lysate with MetaC and palmitate treatment compared to euglycemia (**FIG 3D**).

**Figure 3:**
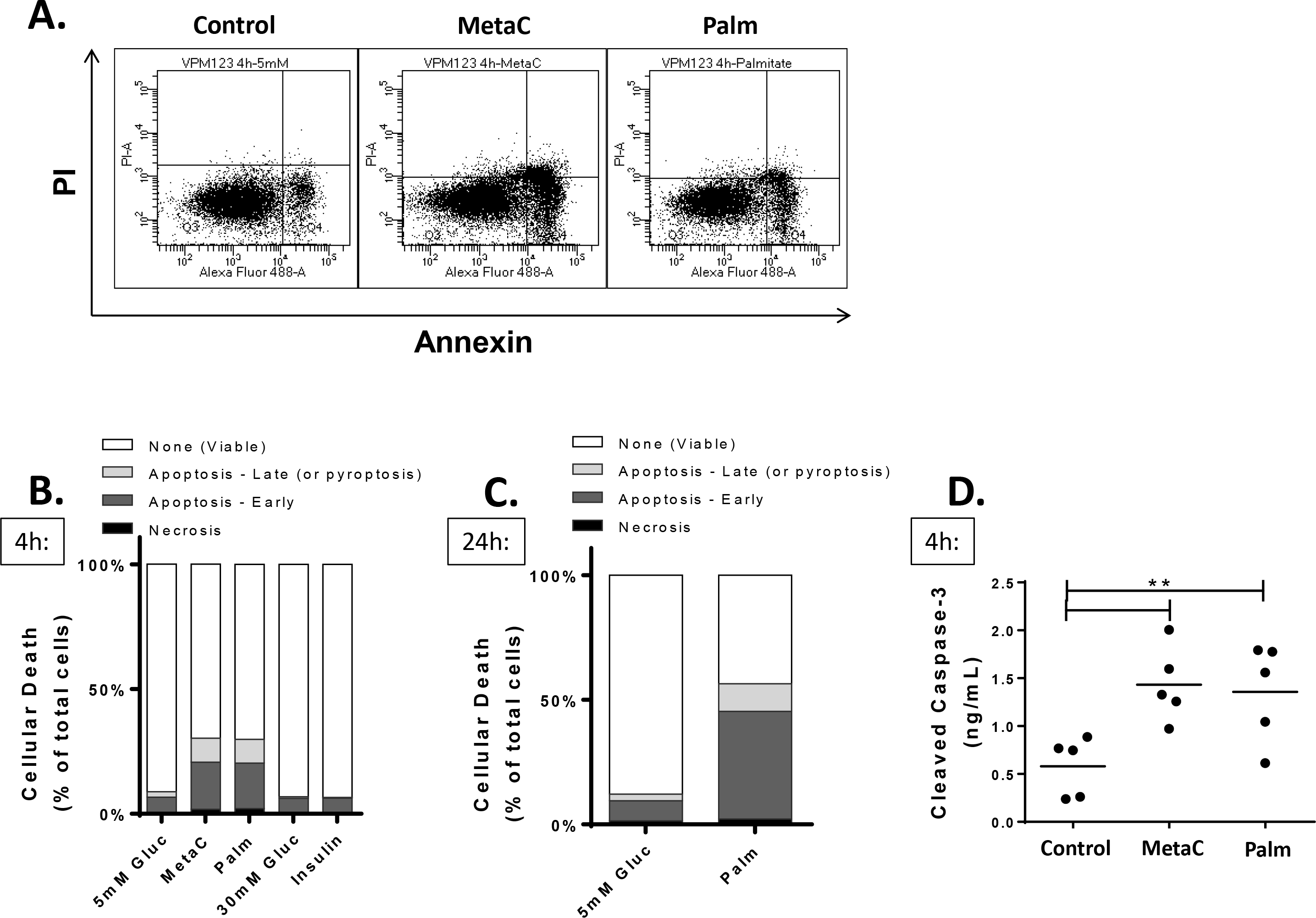
Metabolic stress increases cellular death and cleaves caspase-3 in placental macrophages. Placental macrophages (PMs) were treated for 4 or 24 hours with 5 mM glucose (control), MetaC, palmitate, 30 mM glucose, or insulin as stated in the *Methods* section. Cells were harvested for flow cytometry analysis for cellular death using propidium iodide (PI) and annexin V as markers for necrosis and apoptosis (late & early). (**A)** Representative gating plots for flow cytometry analysis. **(B)** Stacked graph of cellular death averages at 4 h (n=3-14). **(C)** stacked graph of cellular death averages at 24 h (n=5). **(D)** PMs were treated with 5 mM glucose, MetaC, or palmitate (0.4 mM) for 4 h before being lysed and assayed for cleaved caspase-3 by ELISA (n=5). 1-way ANOVA followed by Tukey post-test; mean ± SEM; \*\**p<0.01*.

### Metabolic stress inhibits placental macrophage gene expression of IL-1β

In some macrophages palmitate has been shown to increase IL-1β gene expression [28]. Thus, in response to observing increased IL-1β in the macrophage supernatants following exposure to MetaC or palmitate alone, human placental macrophages were treated with 5 mM glucose (euglycemia) or MetaC for 4 hours and quantitative real-time PCR analysis was performed for IL-1β gene expression. Data show that exposure of placental macrophages to MetaC treatment resulted in a *decrease* in IL-1β mRNA transcript at 4 hours when compared to euglycemia, despite the increased accumulation of IL-1β in the cell supernatant (**SUPPL FIG 5A**).

## Discussion

In the United States and globally, a rising prevalence of noncommunicable diseases, including obesity, type 2 diabetes, GDM and cardiovascular disease, is overburdening healthcare infrastructures and limiting both quality and duration of life [29, 30]. The development of many common non-communicable diseases, such as obesity, diabetes mellitus, cardiovascular disease, and neurocognitive disorders begins years before they are clinically evident in adults. In fact, this risk appears, in part, to be programmed during fetal development in utero, a paradigm known as the Development Origins of Health and Disease [31–33]. Inherent genetic potential combined with environmental factors, such as maternal health and nutrition, influence fetal development [34]. This framework helps explain how parental factors influence health outcomes of offspring and contribute to the transgenerational inheritance of non-communicable disease risk beyond the influence of genetics. The maternal (*in utero*) environment is increasingly recognized as a critical stage for imprinting non-communicable disease risk on the developing fetus [33]. Herein we used *in vitro* models to define direct effects of the metabolic syndrome and hyperglycemia on placental cellular immune responses, which could impact fetal development and the health and wellness of offspring.

Specifically, we interrogated the impact of high ambient levels of glucose, insulin and palmitate on human placental macrophage viability and inflammatory response, finding that only the saturated fat altered the particular readouts we assessed. The primary finding here is that palmitate resulted in the triggering of apoptotic cell death of placental macrophages and induced the NLRP3 inflammasome to become active and release the proinflammatory cytokine IL-1β.

The effects of maternal obesity on placental biology are complex. Maternal meta-inflammation associated with obesity (for example, elevated IL-6, TNF-α, leptin and low adiponectin) may stimulate placental nutrient transport, contributing to fetal overgrowth [35]. Similar inflammatory and hormonal changes may contribute to neurological consequences in offspring via effects on placental serotonin production, while effects on early pregnancy vascularization may predispose some obese women to placental insufficiency, resulting in hypertensive disorders of pregnancy and (seemingly paradoxically) intrauterine growth restriction [35]. Studies of the human placental transcriptome suggest that prepregnancy obesity is associated with significant alterations in the expression of genes involved in lipid metabolism, angiogenesis, hormone activity, and inflammation that corresponded with increased placental lipid content, oxidative stress, and signs of enhanced inflammation [36]. Our data, albeit focused on one particular immune cell from within this complicated heterocellular tissue, support the latter concept regarding inflammation.

The macrophage data presented here support studies by other investigators demonstrating that the human trophoblast responds to saturated fat by activating the NLRP3 inflammasome and releasing IL-β via caspase 1 [16]. This suggests a shared response among immune and parenchymal cells, which could have a biological amplifying effect.

Our studies have limitations that warrant discussion. Here we used placental macrophages, obtained from the villous core of placental cotyledons. However, we did not use genetic or other tests to assess whether these cells were maternal or fetal in origin, though our suspicion based on the isolation technique is that we obtained primarily fetal macrophages (Hofbauer cells). We did not know the prepregnancy body mass index (BMI) of the mothers in this study or other potentially relevant clinical characteristics (such as medication or tobacco exposure histories), which could confound our data and contribute to the intersubject variability observed. We also did not stratify results of these experiments by sex of the offspring, since this was not determined for these studies. This deserves closer attention in future studies, since metabolic stress during pregnancy might exhibit sexually dimorphic results in the placenta [15]. Our results using annexin V and PI staining suggest that palmitate strongly induced apoptosis (annexin+/PI−) in placental macrophages, though we cannot rule out a small contribution of pyroptosis in the dual-staining (annexin+/PI+) cell populations, which we suspect reflect cells stained late after apoptosis onset. Another limitation of our studies is the focus on a prespecified set of macrophage responses to metabolic stress, including particular inflammatory cytokines, caspases, and types of cell death. It is possible that other biological responses were perturbed by our experimental conditions which we might have missed. Future studies will need to employ bias-reduced methods such as transcriptomic, proteomic, or lipidomic analyses.

In summary, we report here that saturated fat induces inflammatory responses in human placental macrophages while also provoking cell death primarily via apoptosis. Future studies will need to determine the significance of these findings *in vivo* and the extent to which maternal covariates such as BMI or fetal covariates such as sex influence macrophage immune responses to saturated fat. These results could be important for understanding how maternal comorbidities impact fetal wellbeing and the health of offspring exposed *in utero* to metabolic stressors.

## Abbreviations

GDM: gestational diabetes mellitus
NCD: non-communicable disease
NLRP3: NLR Family Pyrin Domain Containing 3
FFA: free-fatty acid

## Acknowledgments

We would like to thank the Vanderbilt Institute for Clinical and Translational Research (VICTR, supported by a Clinical and Translational Science Award from the National Center for Advancing Translational Sciences at the National Institutes of Health), for their assistance in this work. The authors have no conflicts of interest to report.

## Funding

This work utilized the Pilot and Feasibility Program of the Vanderbilt Diabetes Research and Training Center funded by grant DK020593 from the National Institute of Diabetes and Digestive and Kidney Disease. AJE was supported by the Childhood Infections Research Program (T32AI095202-02).

---

**Supplemental Figure 1.**
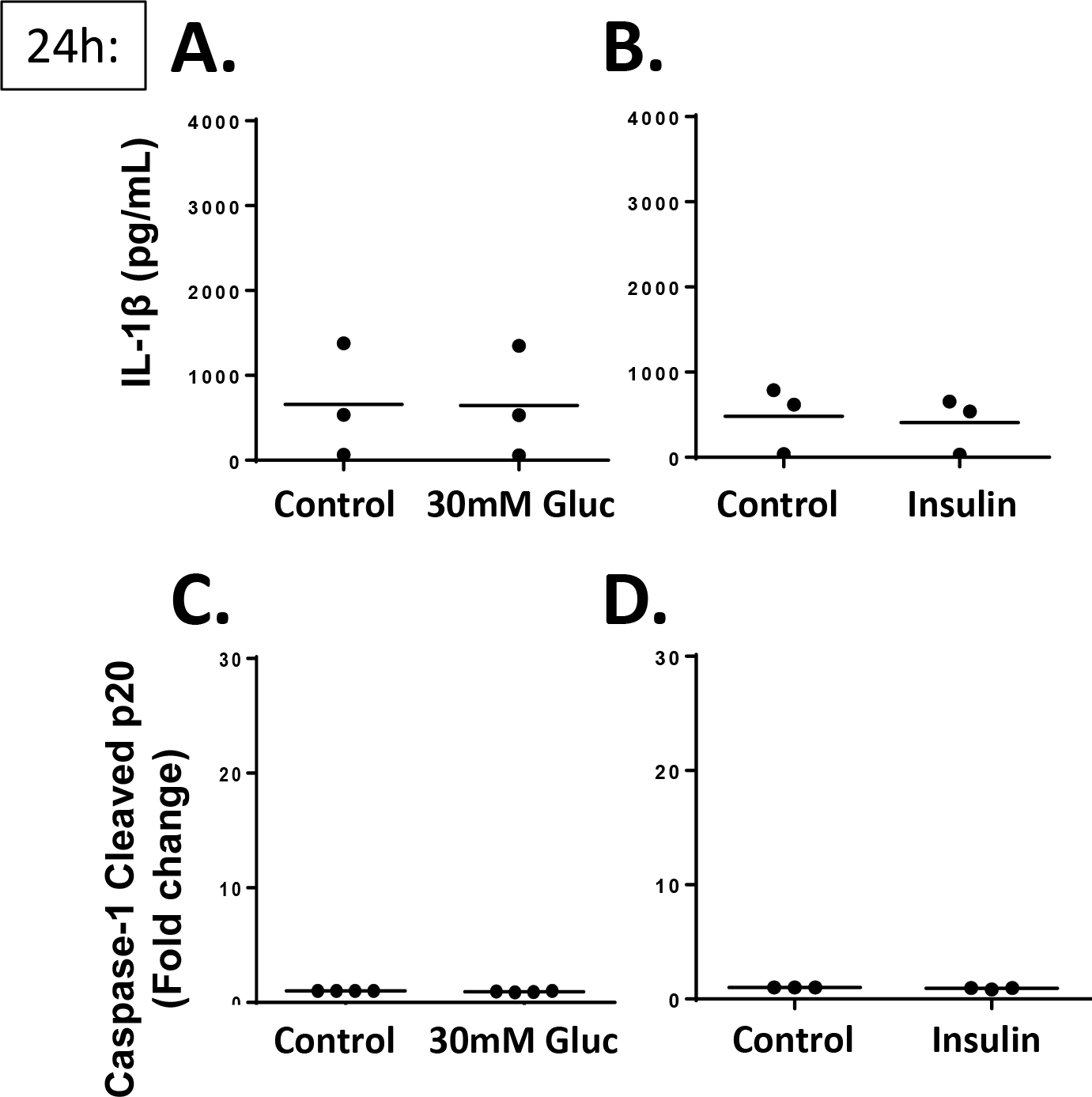
The metabolic stress-induced increase in the release of IL-1β and cleaved caspase-1 in placental macrophages is not due to hyperglycemia or hyperinsulinemia. Placental macrophages were treated for 24 h with 5 mM glucose (control), 30 mM glucose, or insulin (10 nM) as stated in the *Methods* section before supernatant was harvested for IL-1β **(A-B)** (n=3) and cleaved caspase-1 **(C-D)** (n=3-4) ELISA analysis. Paired Student *t*-test; mean ± SEM for all graphs.

**Supplemental Figure 2.**
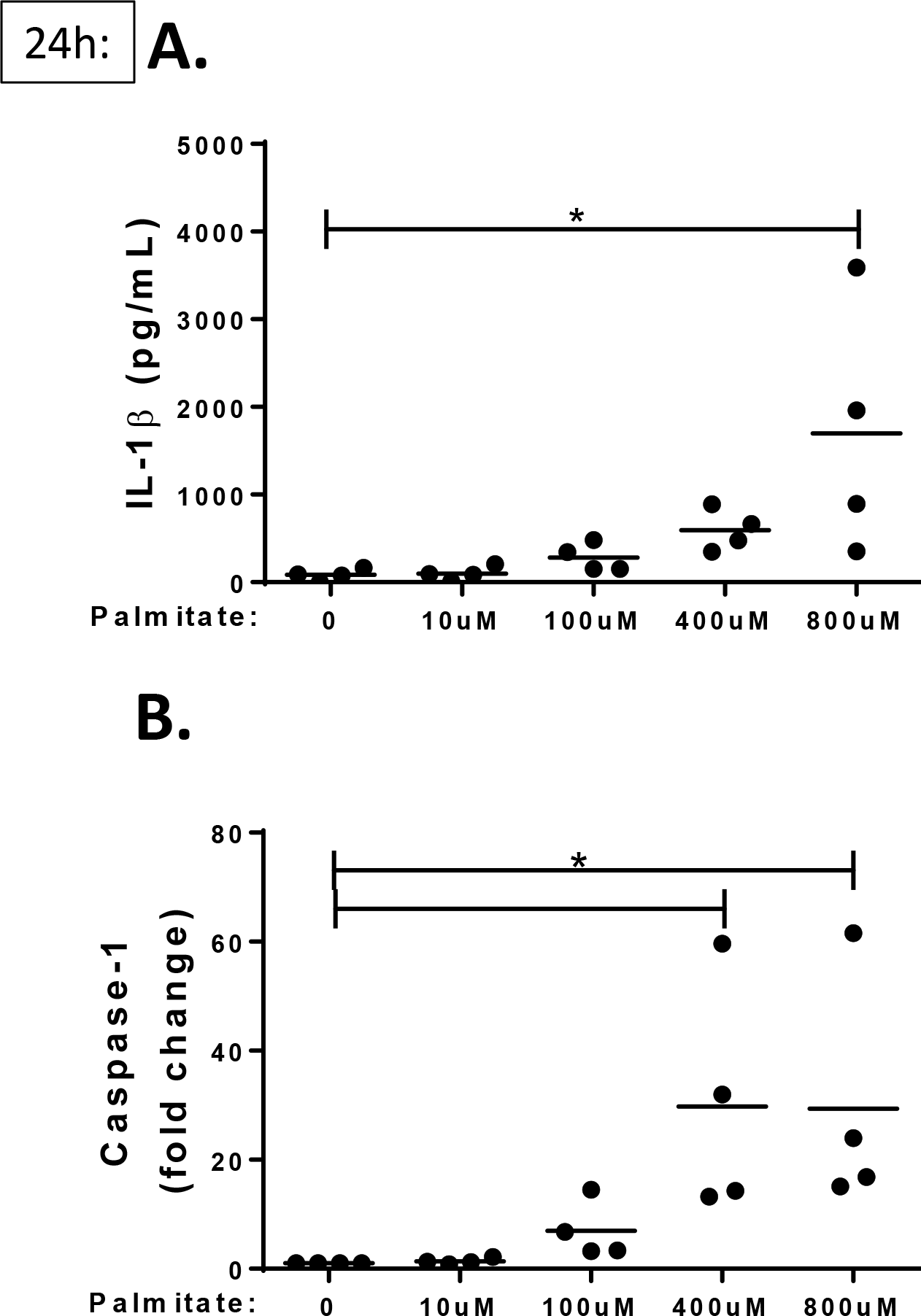
Palmitate increases the release of IL-1β and cleaved caspase-1 in placental macrophages in a dose-dependent manner. Placental macrophages were treated for 24 h with 5 mM glucose (control) or increasing doses of palmitate (0 – 0.8 mM) before supernatant was harvested for IL-1β **(A)** and cleaved caspase-1 **(B)** ELISA analysis (n=4 each). 1-way RM ANOVA with Dunnett post-test; mean ± SEM; *\*p<0.05.*

**Supplemental Figure 3.**
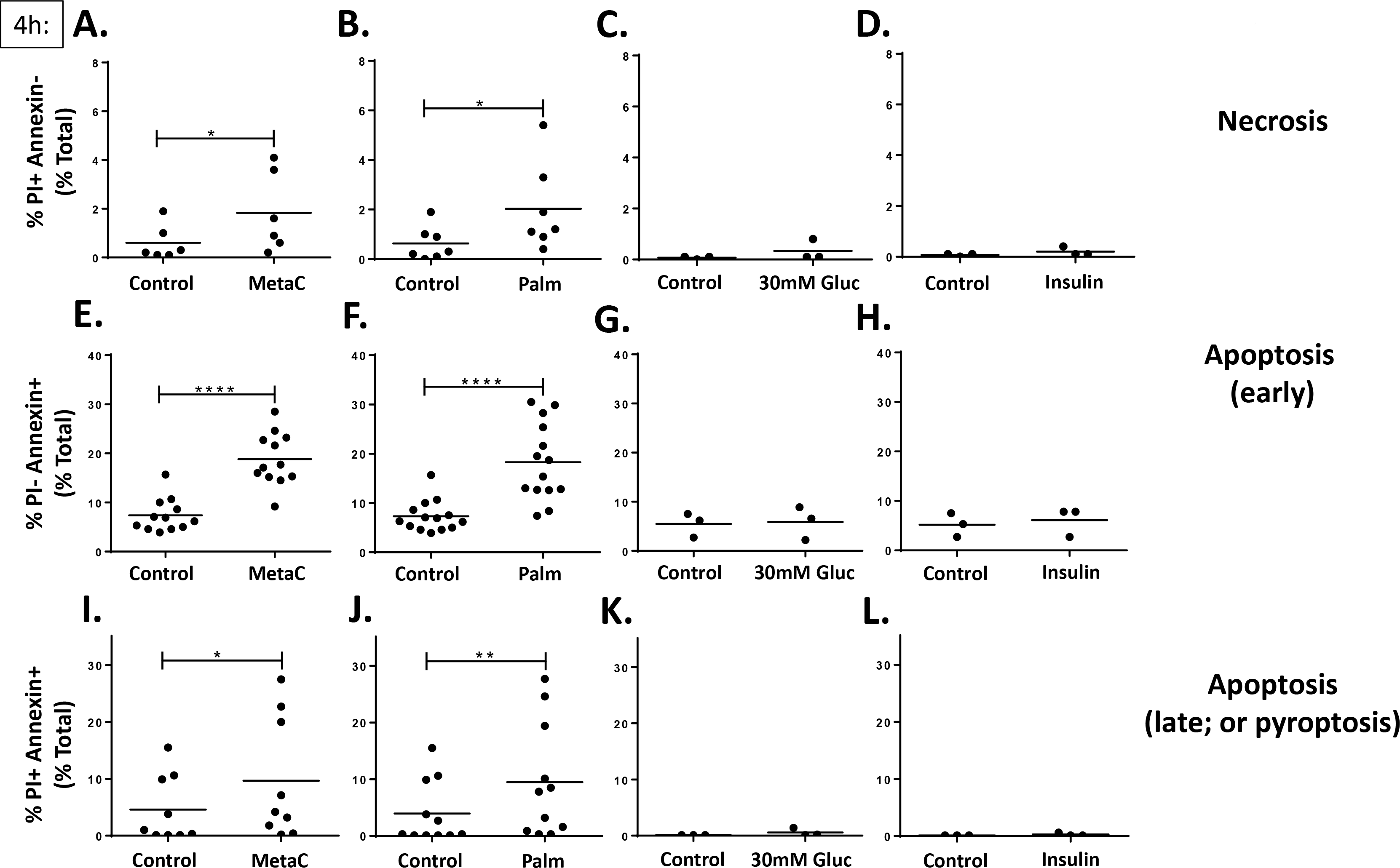
Metabolic stress induces multiple types of cellular death at 4 hours. Placental macrophages were treated for 4 h with 5 mM glucose (control), MetaC, palmitate (0.4 mM), 30 mM glucose, or insulin (10 nM) as stated in the *Methods* section. Cells were harvested for flow cytometry analysis for cellular death using PI and annexin as markers for necrosis and apoptosis (late & early). **(A-D)** PI+annexin− cells defined as necrotic (n=3-7). **(E-H)** PI-annexin+ cells defined as early apoptotic (n=3-14). **(I-L)** PI+annexin+ cells defined as late apoptotic (n=3-11). Paired Student *t*-test; mean ± SEM; \**p<0.05*, \*\**p<0.01*, and \*\*\*\**p<0.0001*.

**Supplemental Figure 4.**
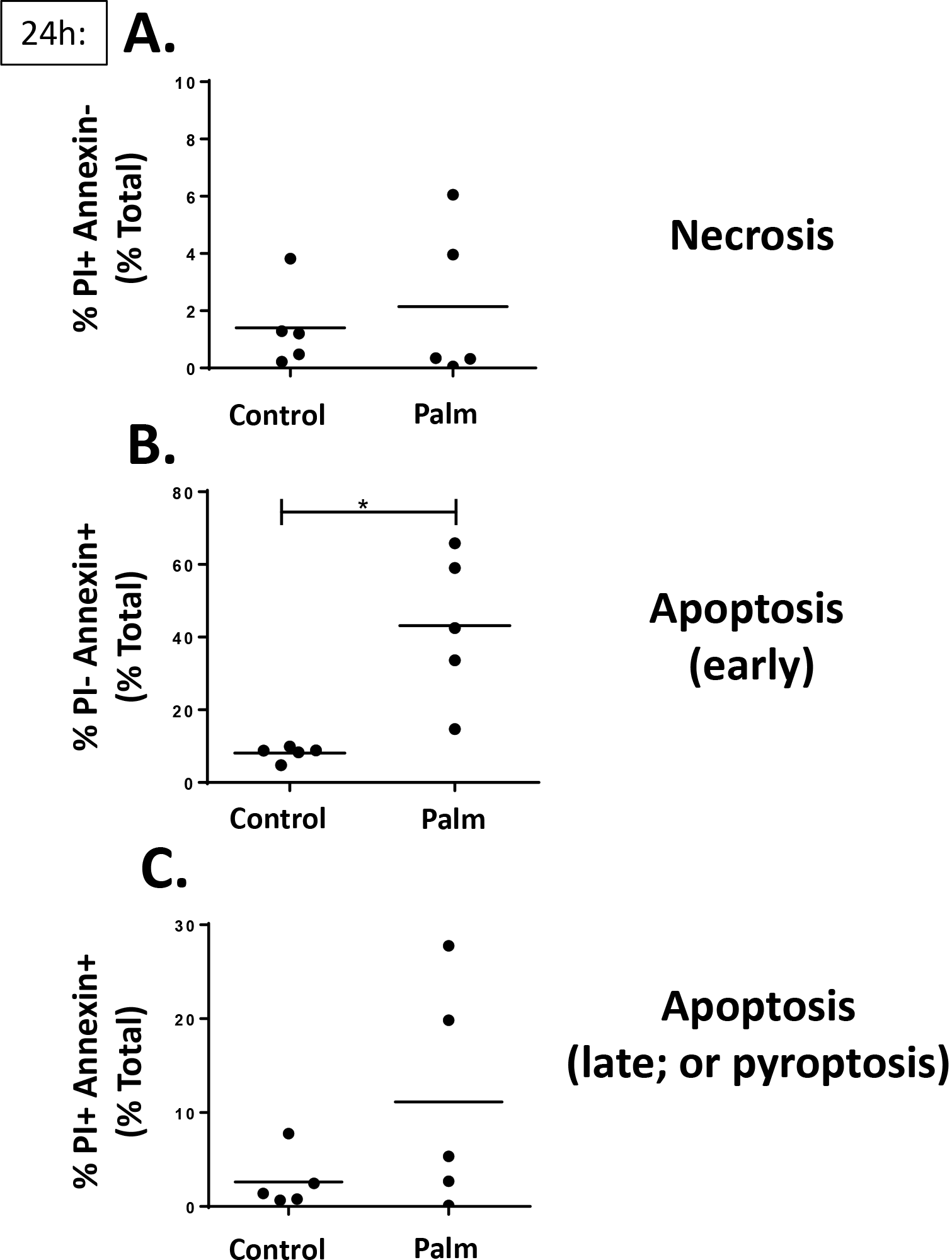
Metabolic stress induces multiple types of cellular death at 24 hours. Placental macrophages were treated with 5 mM glucose (control) or palmitate (0.4 mM) for 24 h. Cells were harvested for flow cytometry analysis for cellular death using PI and annexin as markers for necrosis and apoptosis (late & early). **(A)** PI+annexin-cells defined as necrosis (n=5). **(B)** PI-annexin+ cells defined as early apoptosis (n=5). **(C)**PI+annexin+ cells defined as late apoptosis (n=5). Paired Student *t*-test; mean ± SEM; \**p<0.05*.

**Supplemental Figure 5.**
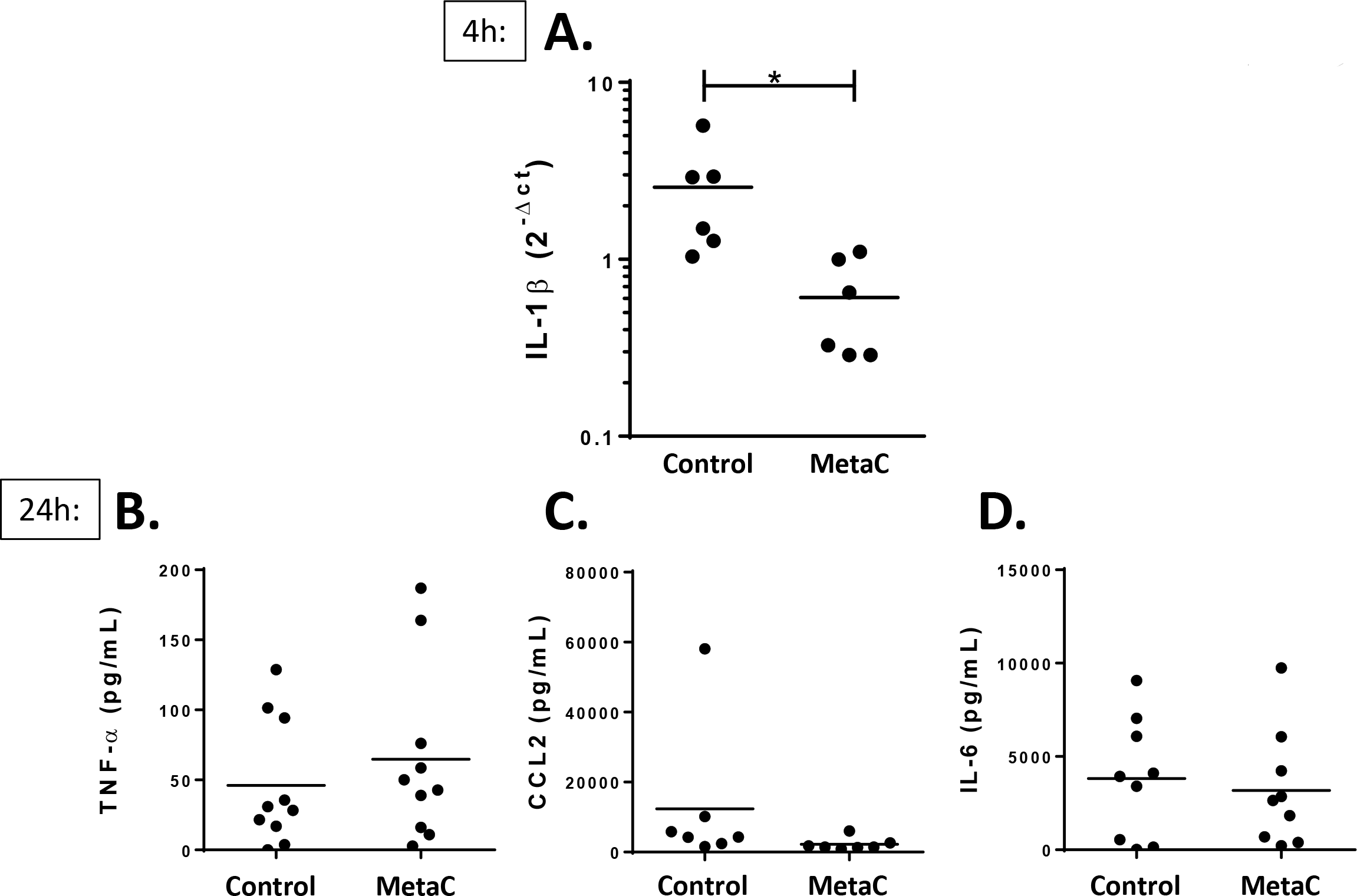
Metabolic stress inhibits gene expression of IL-1β at 4 hours, but did not alter pro-inflammatory cytokine protein levels TNF-α, CCL2, or IL-6 at 24 hours. **(A)** Placental macrophages (PM) were treated with 5 mM glucose (control) or MetaC for 4 h and RNA was harvested for RT-PCR analysis for IL-1β mRNA transcript (n=6). PM cells were treated with 5 mM glucose (control) or MetaC for 24 h and supernatants were harvested for **(B)** TNF-α (n=10), **(C)** CCL2 (n=7), and **(D)** IL-6 (n=9) ELISA analysis with paired Student *t*-tests; mean ± SEM; \**p<0.05*.

